# Overexpression of ELOVL6 has been associated with poor prognosis in patients with head and neck squamous cell carcinoma

**DOI:** 10.1101/2021.02.13.431065

**Authors:** Ruoya Wang

## Abstract

Head and neck squamous cell carcinoma (HNSCC) is a high mortality disease. Extension of long-chain fatty acid family member 6 (ELOVL6) is a key enzyme involved in fat formation that catalyzes the elongation of saturated and monounsaturated fatty acids. Overexpression of ELOVL6 has been associated with obesity-related malignancies, including hepatocellular carcinoma, breast, colon, prostate, and pancreatic cancer. The following study investigated the role of ELOVL6 in HNSCC patients. Gene expression and clinicopathological analysis, enrichment analysis, and immune infiltration analysis were based on the Gene Expression Omnibus (GEO) and the Cancer Genome Atlas (TCGA), with additional bioinformatics analyses. The statistical analysis was conducted in R, and TIMER was used to analyze the immune response of ELOVL6 expression in HNSCC. The expression of ELOVL6 was related to tumor grade. Survival analysis showed that patients with high expression of ELOVL6 had a poor prognosis. Moreover, the results of GSEA enrichment analysis showed that ELOVL6 affects the occurrence of HNSCC through fatty acid metabolism, biosynthesis of unsaturated fatty acids, and other pathways. Finally, ELOVL6 verified by the Human Protein Atlas (HPA) database were consistent with the mRNA levels in HNSCC samples. ELOVL6 is a new biomarker for HNSCC that may be used as a potential predictor of the prognosis of human HNSCC.

## Introduction

Head and neck squamous cell carcinoma (HNSCC) is an aggressive epithelial malignancy worldwide, usually classified in tobacco-related HNSCC and human papillomavirus (HPV)-positive HNSCC. More than 650,000 new cases and 330,000 deaths were reported in 2018(1). Despite technological advances, which have promoted early detection and timely intervention, clinical outcomes and long-term survival rates in HNSCC patients have not improved; the 5-year survival rate remains around 50 percent (2, 3), while the median survival rate of patients with recurrent or metastatic disease is approx.10 months. The high mortality is associated with a high rate of late diagnosis, and the survival rate of late patients is only 34.9 percent(4). Radiotherapy, surgery, and chemotherapy have been considered as the main treatment approaches. Although HNSCC is treated in a variety of ways, patients still have a lower survival probability. Therefore, the discovery of sensitive HNSCC biomarkers remains crucial.

Extension of long-chain fatty acid family member 6 (ELOVL6) is part of a highly conserved endoplasmic reticulum family involved in long-chain fatty acid formation. ELOVL6 catalyzes the elongation of saturated and monounsaturated fatty acids with 12, 14, and 16 carbon atoms to 18 carbon fatty acids (5). ELOVL6 is commonly expressed in high-fat tissues, such as the liver, brown and white adipose tissue, and the brain (6, 7). ELOVL6 up-regulation is involved in insulin resistance. Moreover, ELOVL6 has been associated with obesity-related malignancies, including hepatocellular carcinoma (5), breast, colon, prostate, and pancreatic cancer (8–10). Yet, its role in HNSCC progression remains unclear.

In this work, we used microarray data obtained from the TCGA and GEO database to investigate the expression of ELOVL6 in HNSCC samples. We used R (3.6.3 version) to examine the association of ELOVL6 expression with certain clinical parameters and the prognosis of patients with HNSCC. To better understand the biological processes associated with ELOVL6 regulatory networks, which may be the basis for head and neck squamous cell carcinogenesis, we performed gene set enrichment analysis (GSEA) and Kyoto Encyclopedia of Genes and Genomes (KEGG) analysis. We also used TIMER to detect the association of ELOVL6 with tumor-infiltrating immune cells (TIICs). Besides, we analyzed the correlation between ELOVL6 and HNSCC and the role of ELOVL6 in the occurrence and development of HNSCC.

## Material and methods

### Evidence from TCGA database

Data on gene expression were obtained from the GEO (https://www.ncbi.nlm.nih.gov/gds). Moreover, additional data on gene expression, immune system infiltration (workflow type: HTSeqFPKM), and related clinical information (Data type: Clinical Supplement) were obtained from the TCGA database of HNSCC (https://portal.gdc.cancer.gov/). The clinical factors included gender, stage, age, grade, T-phase, M-phase, N-phase, survival status, and a number of days of survival. We retained RNA-Seq and clinical data for further studies. In addition, R (3.6.3 version) and R BioConductor software packages were used for data analysis. Our study was in accordance with the publication guidelines provided by TCGA.

### Gene enrichment analysis

GSEA was performed using normalized RNA-Seq data obtained from TCGA(11). The number of permutations was set to 1,000. KEGG pathways were analyzed using GSEA to investigate possible biological functions of ELOVL6. Enrichment results had to satisfy one condition, a nominal p-value<0.05. The GSEA created a list of all gene permutations associated with ELOVL6 expression. The samples were then divided into high ELOVL6 groups and low ELOVL6 groups as training sets to distinguish potential functions; GSEA was used to elucidate obvious survival differences. Multiple genome substitutions were performed for each test by the degree of ELOVL6 expression as phenotypic markers. Normalized enrichment scores (NES) and nominal P-values were used to classify the enrichment pathways in each phenotype.

### Immune infiltrates analysis

The potential relationship between ELOVL6 expression and TIICs was evaluated using timer-related modules. TIMER is a comprehensive resource for systematic analysis of immune infiltration in different cancer types (https://cistrome.shinyapps.io/timer/)(12), which relies on a recently published statistical method, deconvolution, to infer TIIC prevalence from gene expression profiles (13). To approximate TIIC abundance, the TIMER database used TCGA data from 10,897 samples of 32 cancers. We examined the expression of ELOVL6 in HNSCC and its relevance to the abundance of TIICs (including B cells, CD8+ T cells, CD4+ T cells, macrophages, neutrophils, and dendritic cells) through gene modules. TIMER produced a graph illustrating gene expression levels against tumor purity(14).

### Data validation

The human protein atlas database (HPA) (www.proteinatlas.org) was used to analyze the protein expression of ELOVL6 between normal and head and neck squamous cell carcinoma tissues. The HPA provides access to 32 human tissues and their protein expression profiles and uses antibody profiling to accurately assess protein localization. Additionally, the HPA provides measurements of RNA levels(15).

### Statistical analysis

All statistical analyses were carried out using R (version 3.6.3). To calculate 95% CI and HR, we used the univariate and multivariate models for Cox analysis. Logistic regression, Wilcoxon rank-sum test, and Kruskal test were used to analyze the correlation between clinical features and ELOVL6 expression. Single-factor survival analysis was used to compare the relationship between several clinical characteristics and survival rates. Using multivariate Cox analysis, we assessed the expression of ELOVL6 and the effects of other pathological and clinical factors (sex, age, grade, lymph node, distant metastasis, tumor status, and stage) on overall survival (OS). The P-value>0.05 expressed by ELOVL6 was set as the threshold.

## Results

### Analysis of survival outcomes and variables

Four HNSCC-related gene expression profiling datasets, including GSE30784, GSE23036, GSE33205, and GSE59102, were obtained directly from the GEO website. GSE30784 consists of 44 normal tissues, 167 cancer tissue samples, and 17 dysplasia tissue samples, among which 17 samples of abnormal tissues were removed. The data were generated using the GPL570 platform. GSE23036 consists of five normal tissues and 63 cancer tissue samples using the GPL571 platform; GSE33205 consists of 25 normal tissue samples and 44 cancer tissue samples. The data were generated using the GPL05175-3188 platform. GSE59102 consists of 13 adjacent tissues and 29 cancer tissue samples using the GPL6480-9577 platform.

We divided the four data sets into two groups: GSE30784 was included in the first group, GSE23036, GSE33205, and GSE59102 in the second group. The second data group was integrated after batch correction.

The results of the two groups were consistent. A significant difference in ELOVL6 expression (P<0.01) was found between normal, and tumor tissues; ELOVL6 was highly expressed in tumor tissue (Fig 1A, B).

**Fig. 1.**
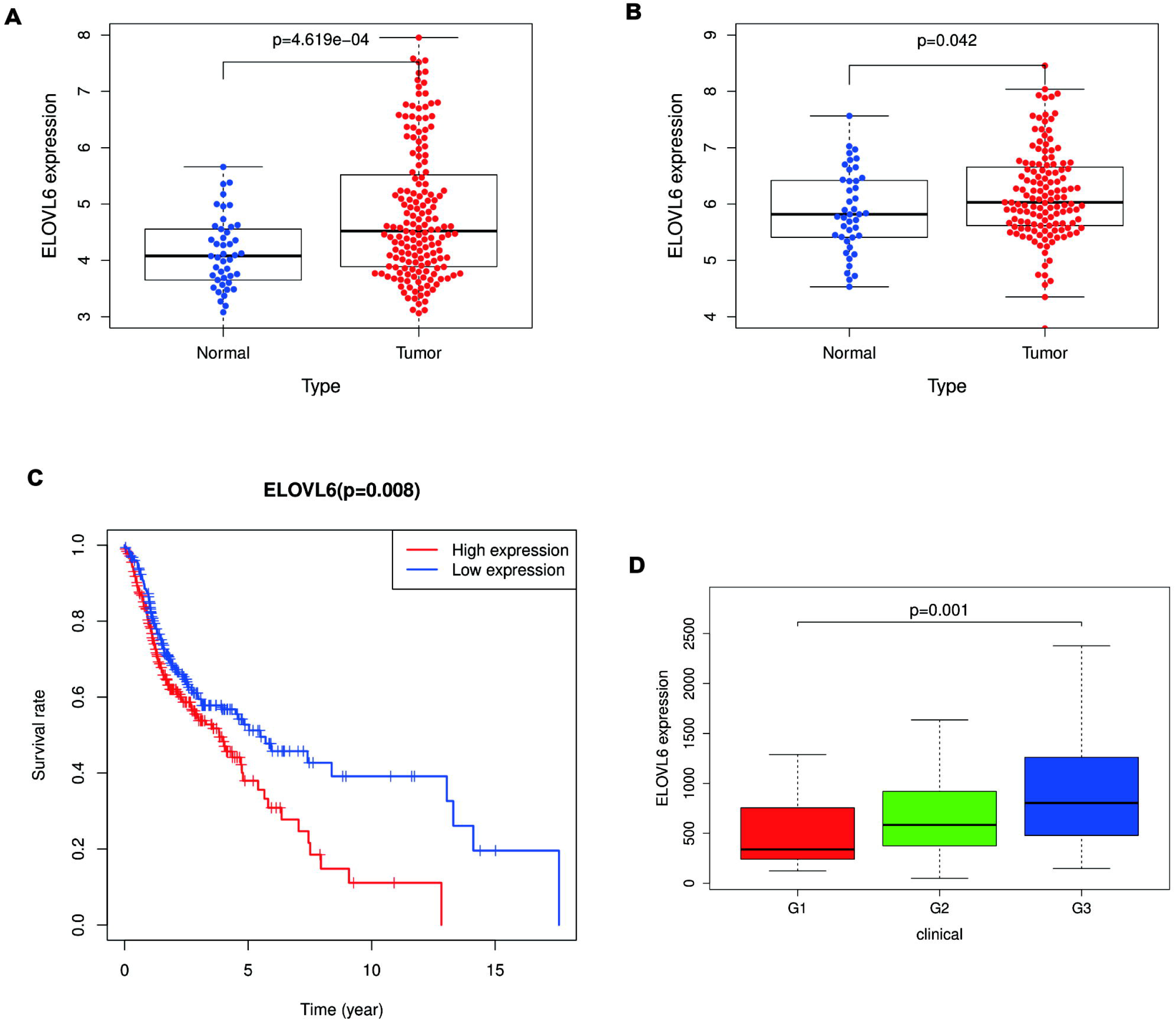
ELOVL6 expression and the association among clinicopathologic factors. (A) The scatter plot showed the difference of ELOVL6 expression between normal and tumor samples in GSE30784 (P<0.01). (B) The scatter plot showing the difference of ELOVL6 expression between normal and tumor samples in GSE23036, GSE33205 and GSE59102 (P<0.01); (C) Survival Analysis of ELOVL6 High and Low Expressions in TCGA (P < 0.01); (D) Expression of ELOVL6 correlated significantly with histological grade in TCGA (P < 0.01).

We then selected 502 tumor samples of HNSCC in the TCGA database. First, we converted the file to alter the count data to more similar values to those obtained from the microarray. Survival analysis indicated that HNSCC with high ELOVL6 expression had a worse prognosis than a tumor with low expression of ELOVL6 (P<0.0; Fig 1C).

Next, we assessed the association between ELOVL6 expression levels with various clinicopathological parameters in patients with HNSCC. The expression of ELOVL6 was found to be significantly correlated with the tumor tissue grade (P<0.01; Fig 1D).

Univariate Logistic regression analysis showed that the expression of ELOVL6 as a well-defined ward variable was related to clinicopathological factors with poor prognosis. The expression of ELOVL6 in HNSCC was significantly correlated with grade (OR=4.4, 95%CI 1.65~12.41, G1 vs. G3). These results suggest that HNSCC patients with high ELOVL6 expression are more likely to develop high-grade tumor (Table 1).

**Table 1.** ELOVL6 expression associated with clinical-pathological characteristics (logistic regression)

| Clinical characteristics | Total(N) | Odds ratio in CENPM expression | P-value |
| --- | --- | --- | --- |
| age (> 65 vs. ≤ 65) | 499 | 0.78(0.53-1.12) | 0.187 |
| Gender (female vs. male) | 500 | 1.10(0.74-1.64) | 0.612 |
| Grade (G1 vs. G2) | 171 | 2.11(0.92-5.09) | 0.08 |
| Grade (G1 vs. G3) | 171 | 4.4(1.65-12.41) | 0.003* |
| Stage (I vs. IV) | 432 | 1.68 ( 0.73-4.00 ) | 0.22 |

The categorical dependent variable, greater or less than the median expression level

### Relationship between ELOVL6 expression and clinicopathology

Cox analysis was used to explore the relationship between ELOVL6 expression and OS and other variable characteristics in patients with HNSCC. Single-factor correlation analysis indicated that stage (HR =1.904, p<0.01), T-phase (HR =1.499, p<0.01), M-phase (HR =20.531, p<0.01), N-phase (HR =1.795, p<0.001), and ELOVL6 mRNA expression (HR=1.457, p<0.01) were significantly correlated with OS. Our multivariate analysis (Fig 2) revealed that ELOVL6 expression (HR=1.39, P=0.015), T-phase (HR=1.70, P=0.017), M-phase (HR=13.16, P=0.021), N-phase (HR=1.93, P=0.000) were independent prognostic factors in patients with HNSCC (Table 2).

**Fig. 2.**
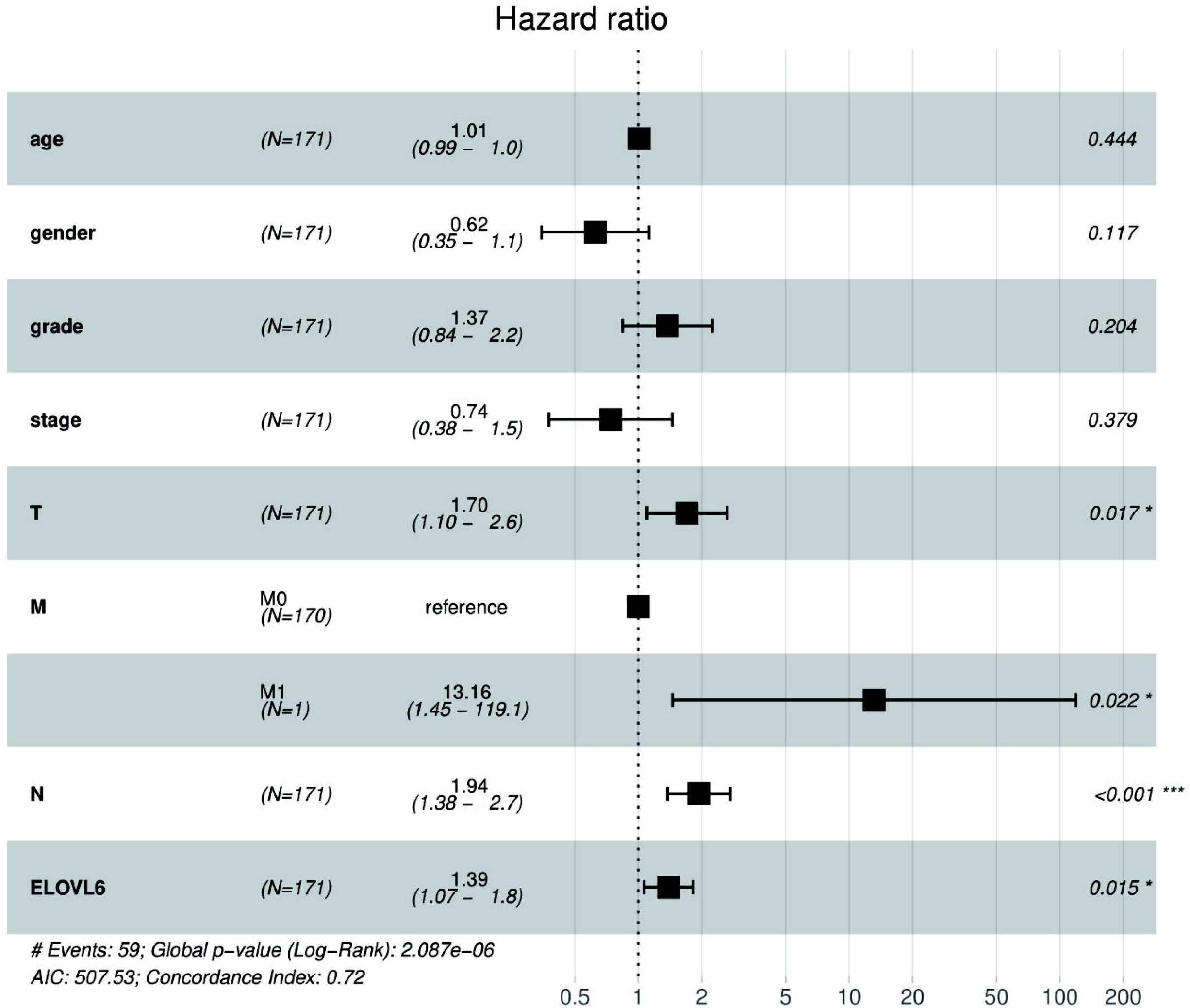
Multivariate Cox analysis of ELOVL6 expression and other clinicopathological variables.

**Table 2.** Correlation between overall survival and multivariable characteristics in TCGA patients via Cox regression and Multivariate survival model.

| Parameter | Univariate analysis |  |  | Multivariate analysis |  |  |
| --- | --- | --- | --- | --- | --- | --- |
|  | HR | 95%CI | P | HR | 95%CI | P |
| Age | 1.01 | 0.98-1.03 | 0.286 | 1.00 | 0.98-1.03 | 0.443 |
| Gender | 0.64 | 0.36-1.14 | 0.133 | 0.62 | 0.34-1.12 | 0.116 |
| Grade | 1.47 | 0.98-2.22 | 0.062 | 1.37 | 0.84-2.24 | 0.203 |
| Stage | 1.90 | 1.23-2.92 | 0.003** | 0.73 | 0.37-1.45 | 0.379 |
| T | 1.49 | 1.13-1.98 | 0.004** | 1.70 | 1.09-2.63 | 0.017* |
| M | 20.53 | 2.56-164.16 | 0.004** | 13.16 | 1.45-119.14 | 0.021* |
| N | 1.79 | 1.34-2.39 | 7.19E-05*** | 1.93 | 1.37-2.72 | 0.000*** |
| ELOVL6 | 1.45 | 1.12-1.88 | 0.004** | 1.39 | 1.06-1.82 | 0.015* |

P-value significant codes: 0 ≤ \*\*\* < 0.001 ≤ \*\* < 0.01 ≤ \* < 0.05

### Analysis of ELOVL6 using GSEA

GSEA was used to explore the potential biological functions of ELOVL6 through KEGG pathway analysis. Significant differences in the enrichment of high levels of ELOVL6 in KEGG pathways were found (p<0.050). According to the Normalized enrichment score (NES), highly enriched signaling pathways were selected. KEGG pathway enrichment analysis revealed nine categories positively associated with high levels of ELOVL6, as shown in Table 3: fatty acid metabolism, biosynthesis of unsaturated fatty acids, certain cancer, WNT signaling pathway, RNA degradation, cell cycle, insulin signaling pathway, and tight junction. The results showed that ELOVL6 high expression differentially enriched fatty acid metabolism, biosynthesis of unsaturated fatty acids, certain cancer, WNT signaling pathway, RNA degradation, cell cycle, insulin signaling pathway, and tight junction (Fig 3).

**Table 3.**
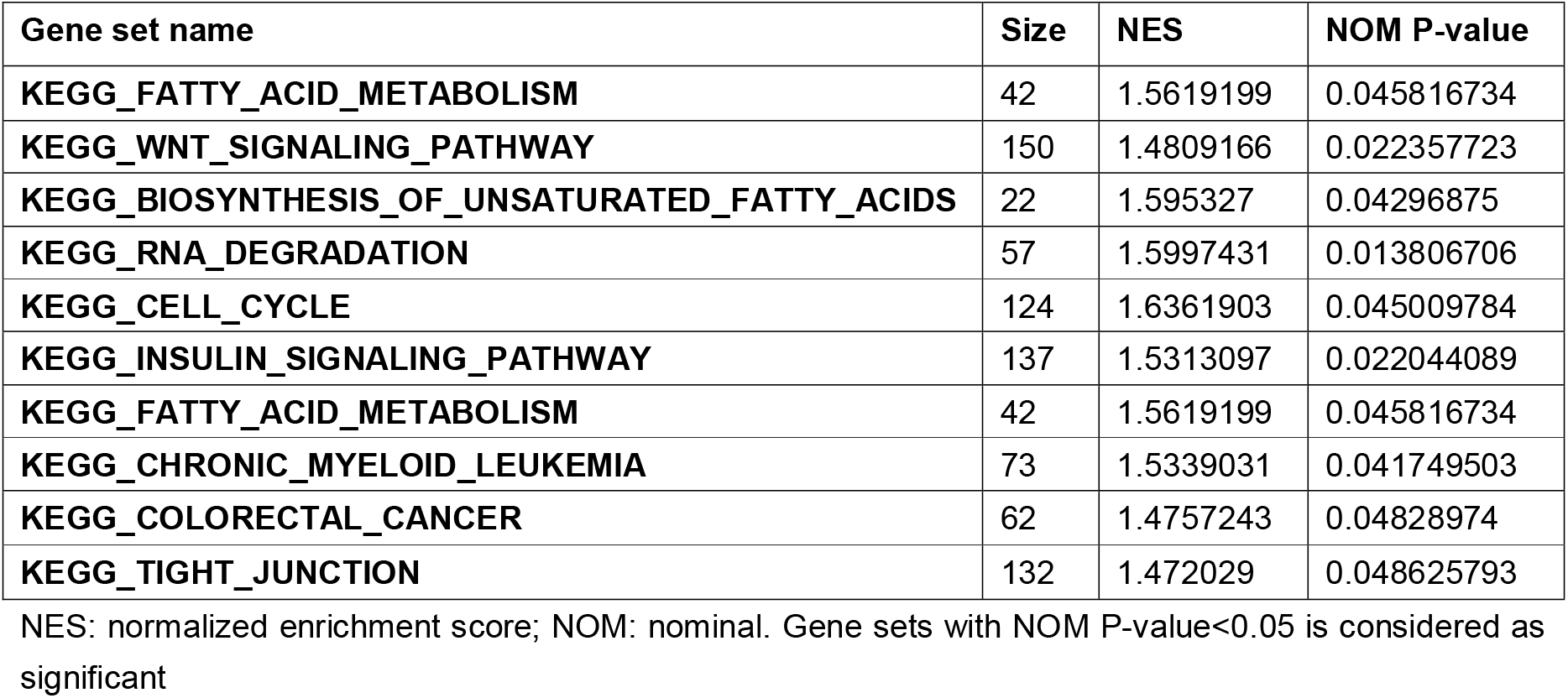
Gene Sets Enriched in Phenotype High.

**Fig. 3.**
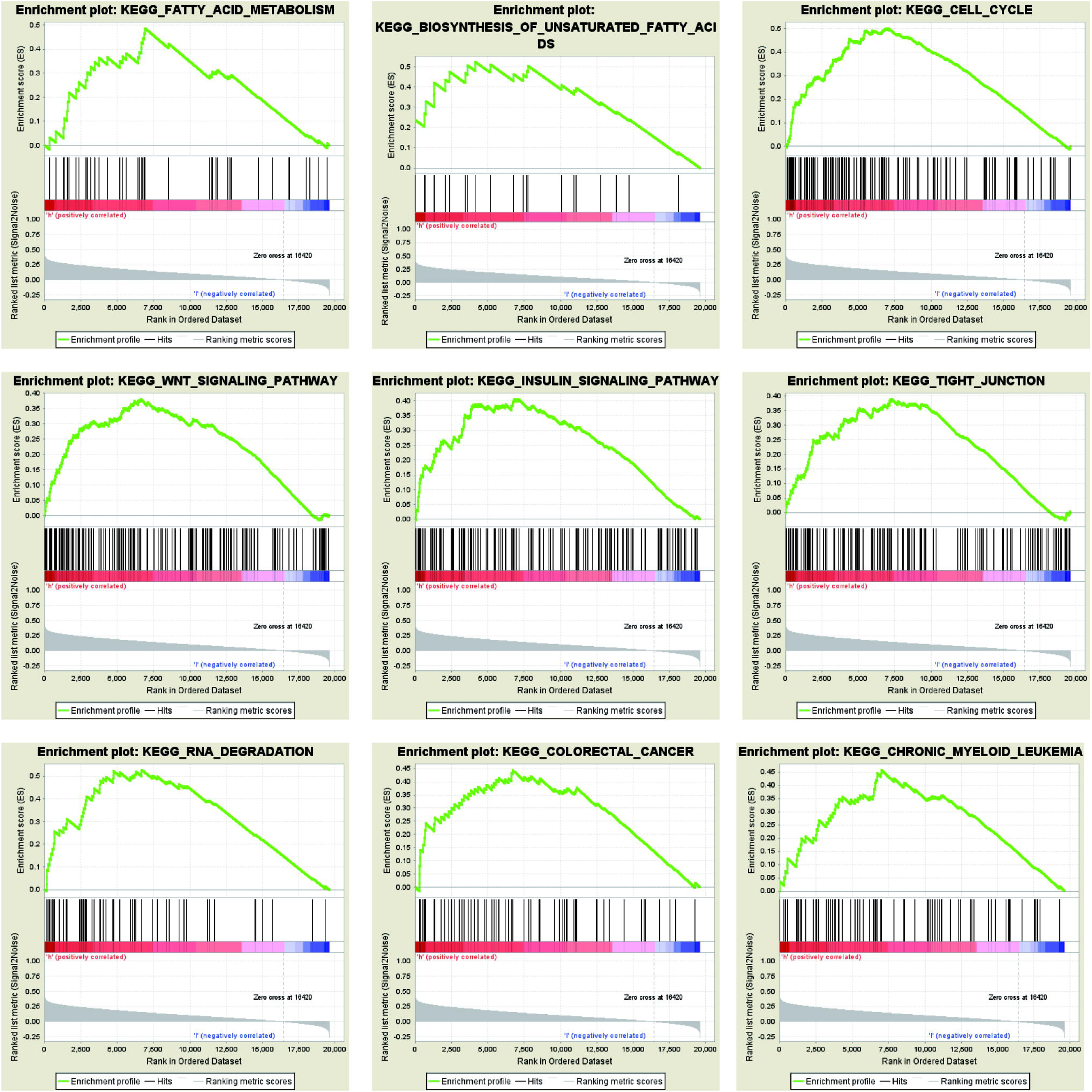
Enrichment plots from gene set enrichment analysis (GSEA)

Independent tumor-infiltrating lymphocytes have an essential role in predicting overall survival and sentinel lymph node status(16). Therefore, in this study, we used TIMER to analyze the possible correlation between ELOVL6 expression and immune infiltration level in HNSCC. ELOVL6 expression was positively correlated with B cells (p=3.72e-01), CD8+ T cells (p=1.48e-01), CD4+ T cells (p=2.92e-02), macrophages (p=1.07e-01), dendritic cells (p=2.43e-01) and negatively correlated with neutrophils (p=5.04e-01), as shown in (Fig 4A). These results suggest that ELOVL6 has a key role in the immune invasion of HNSCC.

**Fig. 4.**
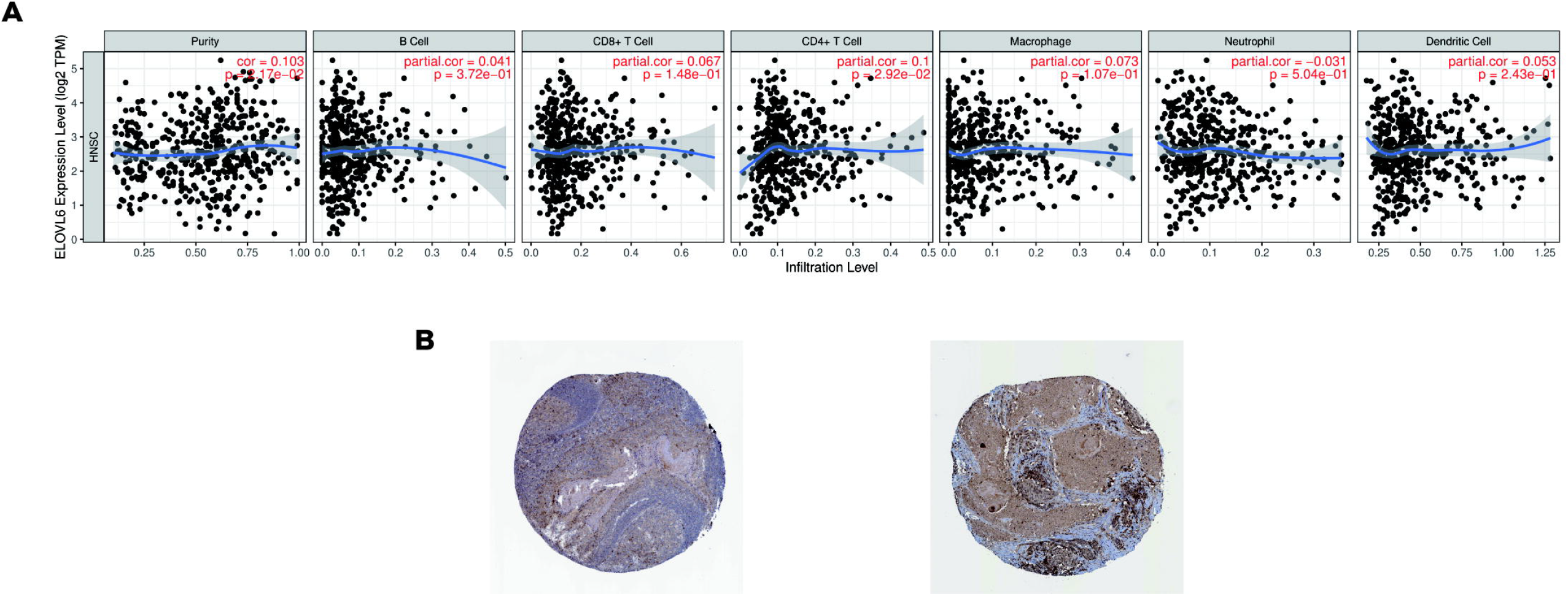
(A) Correlations between ELOVL6 expression and immune infiltration levels. (B) Immunohistochemistry of ELOVL6 based on the Human Protein Atlas.

### Data validation

The ELOVL6 protein levels, verified by the HPA database, were all increased, which was consistent with the mRNA levels in head and neck squamous cell carcinoma samples (Fig 4B).

## Discussion

Extension of long-chain fatty acid family member 6 (ELOVL6) is part of a highly conserved endoplasmic reticulum family involved in long-chain fatty acid formation. ELOVL6 catalyzes the saturation and elongation of monounsaturated fatty acids. Dietary polyunsaturated fatty acids may severely inhibit ELOVL6 expression (6). Moreover, overexpressed ELOVL6 has been found in several cancers, including non-alcoholic steatohepatitis-related hepatocellular carcinoma (17), squamous cell carcinoma (18), and breast cancer (19, 20). Phospholipids containing longer acyl chains are abundant in cancer tissues, and ELOVL6 is the main enzyme responsible for prolonging fatty acids in cancer(18). This elongation has been detected in non-alcoholic steatohepatitis (NASH) associated with hepatocellular carcinoma (17). Moreover, Marien *et al* suggested that inhibition of ELOVL6 might be a potential treatment for lung squamous cell carcinoma(18). In addition, ELOVL6 overexpression has been associated with axillary lymph node metastasis and short disease-free survival in breast cancer(19).

Our results indicate that the expression of ELOVL6 was related to clinicopathological factors (grade), survival time, and poor prognosis in patients with HNSCC. Univariate analysis found that ELOVL6 expression, as a clear-cut variable, was associated with clinicopathological factors with poor prognosis. Stage, T-phase N-phase and M-phase, may have an indispensable role in the further progression of tumors. Univariate and multivariate analysis also showed that ELOVL6 was still closely related to OS. Patients with high expression of ELOVL6 had a decreased survival rate, which was consistent with Martin *et al* (21). To sum up, these data suggest that ELOVL6 may be a potential prognostic biomarker and therapeutic target for the prognosis of HNSCC; however, further studies are warranted.

KEGG pathway analysis indicated that the upregulation of ELOVL6 was mainly related to fatty acid metabolism, WNT signaling pathway, cell cycle, RNA degradation, and then controlled the occurrence and development of cancer cells. More and more studies have shown that cell metabolic disorders are associated with tumor development(22, 23). Metabolic diseases, such as obesity and diabetes are associated with increased risk of cancer, including hepatocellular carcinoma, breast, colon, prostate, and pancreatic cancer(5, 8–10). ELOVL6 is involved in fatty acid metabolic pathways. Increased generation of new fat is an early and common event in the development of cancer(24). Lipogenesis is considered as a potential target for cancer therapy. Some enzymes associated with lipogenesis have been reported as targets for cancer therapy (25).

Upregulation of ELOVL6 expression affects WNT signaling pathways critical to tissue development and homeostasis by regulating their endogenous stem cells. WNT signaling abnormalities can affect the behavior of cancer stem cells (CSC) and trigger and/or maintain and develop many cancers (26).

ELOVL6 upregulation affects tight junctions. Claudins are integral transmembrane proteins in tight junctions essential to maintain cell adhesion and polarity. Changes in individual claudin expression have been detected in cancer and appear to be associated with tumor progression (27).

So far, no association between ELOVL6 and tumor immune response has been reported. In the present study, we used online tools to analyze the correlation between immune infiltration in HNSCC and ELOVL6. TIMER database was applied to analyze the link between ELOVL6 expression and immune infiltration levels in HNSCC. ELOVL6 had the strongest association with B cells, CD8+T cells, CD4+T cells, neutrophils, macrophages, and dendritic cells. HNSCC, especially an oropharyngeal tumor, is a highly immune-infiltrating tumor (28, 29). CD8+ T cells are the main anticancer effector cell subsets in HNSCC. Their function is often hampered by overexpression of immune checkpoint molecules such as programmed cell death protein 1(PD-1), programmed cell death 1 ligand 1(PD-L1), or cytotoxic T lymphocyte-associated protein 4 (CTLA-4)(30). These molecules have become targets for immunotherapy approaches that alter cancer treatment patterns. At the same time, lymphocytes B are another critical component of TILs that are associated with good prognosis in several human cancer types(31). Large numbers of B lymphocytes in lymph node metastasis are associated with better prognosis in HNSCC patients (32). In the tumor microenvironment, dendritic cells through blood circulation and interact with tumor cells in the tumor microenvironment. At different stages of tumor progression, dendritic cells may exhibit different functions, both as immune stimulators and as immunosuppressive factors (33). All these suggest that ELOVL6 may have an important role in tumor immune response and be a good target for immunotherapy. The association with various tumor characteristics and immune cell responses highlights the role of ELOVL6 upregulation as an independent prognostic factor for poor overall survival.

HNSCC patients with higher expression of ELOVL6 are more likely to develop advanced grade tumors compared to those with low expression in patients with HNSCC. ELOVL6 may affect tumorigenesis mechanism and tumor immunological progression in HNSCC, which suggests that ELOVL6 has a vital role in tumor immune response and can be a good target for immunotherapy.

## Conclusions

To the best of our knowledge, this is the first study that confirmed the importance of ELOVL6 in the prognosis of HNSCC. However, future clinical trials are needed to validate these results. With further understanding of its functional scope, ELOVL6 may become a useful tool for diagnosing and treating HNSCC and promoting the application of ELOVL6 in the prognostic evaluation of HNSCC. Moreover, biomarker therapy may be regarded as a promising option for the treatment of HNSCC.

## Acknowledgements

I thank MedSci for language editing the manuscript draft.

